# *Toxoplasma gondii* PROP1 is critical for autophagy and parasite viability during chronic infection

**DOI:** 10.1101/2024.10.02.616283

**Authors:** Pariyamon Thaprawat, Shreya Chalasani, Tracey L. Schultz, Manlio Di Cristina, Vern B. Carruthers

**Affiliations:** Department of Microbiology and Immunology, University of Michigan Medical School, Ann Arbor, MI, USA; Medical Scientist Training Program, University of Michigan Medical School, Ann Arbor, MI, USA; Department of Chemistry, Biology and Biotechnology, University of Perugia, Perugia, Italy

## Abstract

Macroautophagy is an important cellular process involving lysosomal degradation of cytoplasmic components, facilitated by autophagy-related proteins (ATGs). In the protozoan parasite *Toxoplasma gondii*, autophagy has been demonstrated to play a key role in adapting to stress and persistence of chronic infection. Despite limited knowledge about the core autophagy machinery in *T. gondii*, two PROPPIN family proteins (TgPROP1 and TgPROP2) have been identified with homology to Atg18/WIPI. Prior research in acute stage tachyzoites suggests that TgPROP2 is predominantly involved in a non-autophagic function, specifically apicoplast biogenesis, while TgPROP1 may be involved in canonical autophagy. Here, we investigated the distinct roles of TgPROP1 and TgPROP2 in chronic stage *T. gondii* bradyzoites, revealing a critical role for TgPROP1, but not TgPROP2, in bradyzoite autophagy. Conditional knockdown of TgPROP2 did not impair bradyzoite autophagy. In contrast, TgPROP1 KO parasites had impaired autolysosome formation, reduced cyst burdens in chronically infected mice, and decreased viability. Together, our findings clarify the indispensable role of TgPROP1 to *T. gondii* autophagy and chronic infection.

**Importance:** It is estimated that up to a third of the human population is chronically infected with *Toxoplasma gondii*; however, little is known about how this parasite persists long term within its hosts. Autophagy is a self-eating pathway that has recently been shown to play a key role in parasite persistence, yet few proteins that carry out this process during *T. gondii* chronic infection are known. Here, we provide evidence for a non-redundant role of TgPROP1, a protein important in the early steps of the autophagy pathway. Genetic disruption of TgPROP1 resulted in impaired autophagy and chronic infection of mice. Our results reveal a critical role for TgPROP1 in autophagy and underscore the importance of this pathway in parasite persistence.

## Introduction

Macroautophagy (hereafter, autophagy) is an important cellular homeostatic pathway that involves lysosomal-dependent turnover of cellular materials (1, 2). The sequestration of contents into double-membraned autophagosomes requires multiple autophagy-related proteins (ATGs), many of which are conserved among eukaryotes (3, 4). A key early step in the de novo formation of autophagosomes is the generation of phosphatidylinositol-3-phosphate (PtdIns3P) on autophagic membranes that is essential for proper localization and recruitment of the core autophagy machinery. A class III phosphatidylinositol 3-kinase (PtdIns3K) complex containing the key lipid kinases Vps34 and Vps15 is responsible for the production of PtdIns3P upon autophagy initiation (5–7). PtdIns3P present on autophagic membranes recruit protein scaffolds important for autophagic flux, including the Atg2-Atg18 complex that mediates lipid transfer for membrane expansion (8, 9). A family of PROPPINs (β-propeller that bind phosphoinositides) are required for distinct subtypes of autophagy including Atg18, Atg21, and Hsv2 in *Saccharomyces cerevisiae* and WIPIs 1-4 (WD-repeat protein interacting with phosphoinositides) in mammalian cells (10–13). PROPPINs form seven bladed β-propellers through their WD40 domains that facilitate their binding to phospholipids (14).

While much of the autophagy machinery has been discovered and characterized in yeast and mammalian systems, far less is known about the proteins involved in this pathway in early branching, divergent eukaryotes (15–17). *Toxoplasma gondii* is a eukaryotic pathogen that has been shown to rely on autophagy for cellular survival in response to stress (15, 18–21). Despite a seemingly reduced repertoire of autophagy proteins present in *T. gondii*, there is extensive evidence that this protozoan parasite possesses a functional autophagy pathway. Both the acute stage (tachyzoites) and chronic stage (bradyzoites) parasite forms rely on autophagy for survival from extracellular stressors or to maintain chronic infection, respectively (18–21). Specifically, the putative phospholipid scramblase TgATG9 was recently shown to be required for autophagosome biogenesis and bradyzoite survival (19, 20). However, many of the other discovered TgATGs including the ubiquitin-like TgATG8 serve non-autophagy roles in maintenance of a plastid-organelle, the apicoplast, which houses critical metabolic pathways (22–25). Distinguishing between identified TgATGs that are solely involved in autophagy rather than maintenance of the apicoplast is therefore important to further our understanding of the autophagy pathway in *T. gondii*.

Two PROPPINs (TgPROP1 and TgPROP2) have been identified in *T. gondii* based on homology-directed searches for WD40 domain containing proteins with similarities to *S. cerevisiae* Atg18. In tachyzoites, TgPROP1 appears to be important for the parasite stress response, whereas TgPROP2 has been shown to play a role in apicoplast biogenesis (26, 27). Yet interestingly, both TgPROP1 and TgPROP2 were found to localize to autophagic vesicles in a PtdIns3P-dependent manner upon starvation in tachyzoites (26). Hence, the specific contributions of TgPROP1 and TgPROP2 to *T. gondii* autophagy have not been clearly defined. Addressing this gap in knowledge would determine whether *T. gondii* possesses any forms of redundancy in its core autophagy machinery like the partial redundancies observed among the multiple paralogs of PROPPINs found in yeast and mammalian cells (13, 28, 29).

In this study, we sought to determine the individual contributions of TgPROP1 and TgPROP2 to parasite autophagy. We provide evidence that TgPROP2 is not critical for bradyzoite autophagy and reveal new insights into the non-redundant role of TgPROP1.

## Results

### Conditional knockdown of TgPROP2 does not affect bradyzoite autophagy

While it has been reported that TgPROP2 is required for apicoplast biogenesis, the contribution of TgPROP2 to bradyzoite autophagy has not been defined (27). To determine if TgPROP2 is necessary for bradyzoite autophagy, we generated a conditional knockdown mutant of TgPROP2 using the auxin-inducible degron (AID) system since TgPROP2 is essential in tachyzoites (26, 30). More specifically, in cystogenic ME49Δku80Δhxg strain parasites expressing the heterologous F-box protein TIR1 (termed TIR1 hereafter) we endogenously tagged TgPROP2 at its C terminus with a minimal auxin-inducible degron (mAID) fused to mNeonGreen (Figure 1A) (31). Upon addition of auxin (indole-3-acetic acid, IAA), we observed marked knockdown of TgPROP2-mAID in *in vitro* differentiated bradyzoites after 72 h by western blotting and immunofluorescence microscopy (Figure 1B-C). Based on our western blot, IAA-treated bradyzoites had minimal TgPROP2 protein, at least less than 33.3% of the untreated controls (Figure 1B). We assessed the impact of conditional knockdown of TgPROP2-mAID on bradyzoite autophagy by differentiating *in vitro* bradyzoites for 7 days followed by treatment with IAA or vehicle control with or without LHVS for 3 days prior to CytoID staining (18, 19, 32, 33). LHVS is a potent inhibitor of TgCPL and results in the accumulation of undigested autophagic material delivered to the PLVAC (34). Bradyzoites of the TgPROP2-mAID or TIR1 parental strains accumulated CytoID-positive autolysosome puncta in the presence or absence of IAA treatment. As a positive control, we included conditional knockdown of another autophagy protein, TgATG9, which was recently reported to be important in autophagosome biogenesis based on its failure to generate CytoID-positive autolysosomes after IAA treatment (20) (Figure 1D-E). These findings indicate that conditional knockdown of TgPROP2 does not impact autolysosome formation in bradyzoites and that its role might be dedicated to apicoplast biogenesis, congruent with previous reports (26, 27).

**Figure 1.**
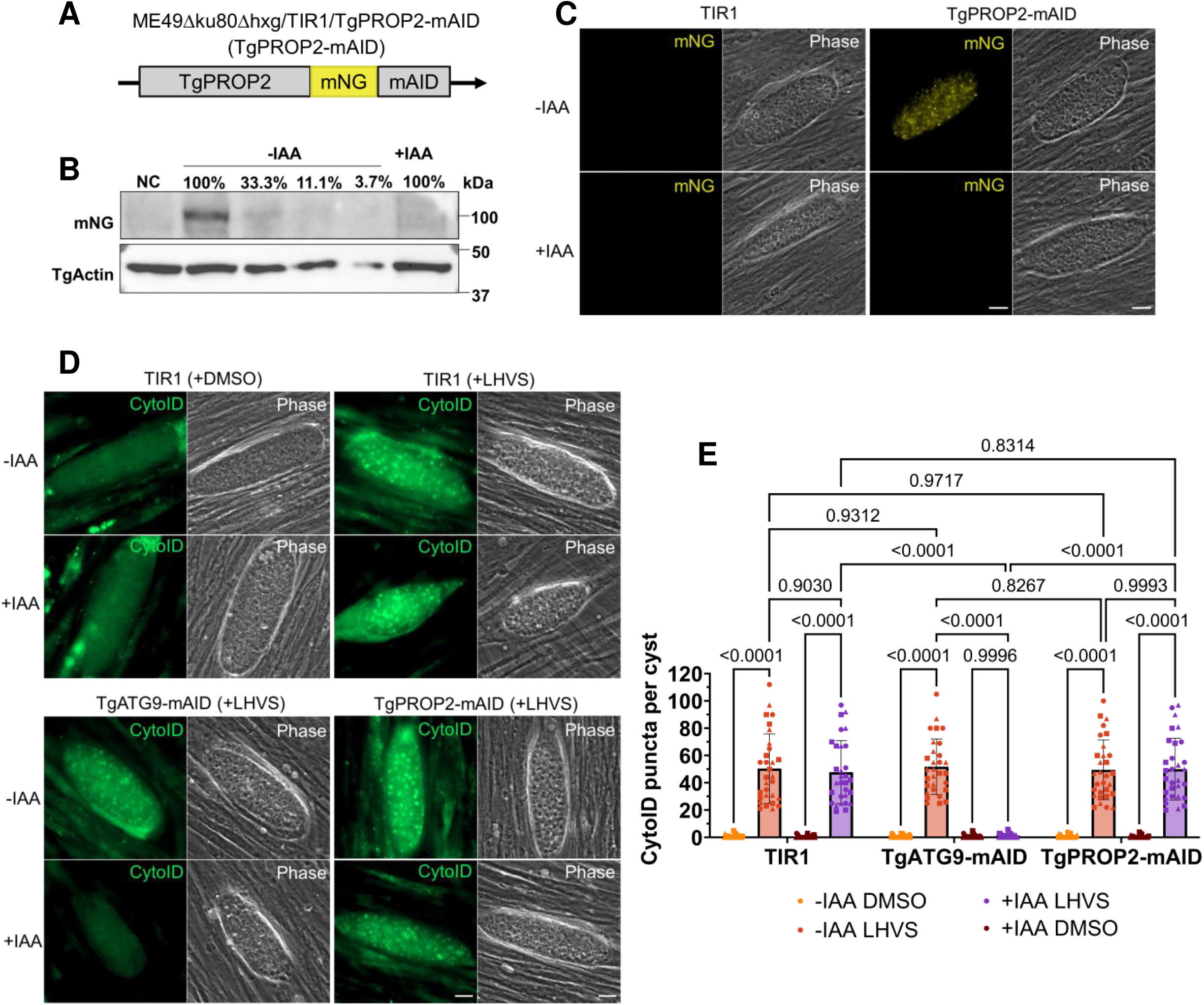
TgPROP2 is not required for bradyzoite autophagy. (**A**) Endogenous tagging of TgPROP2 with mNeonGreen (mNG) and the minimal auxin inducible degron (mAID) was achieved with CRISPR-Cas9 and homology directed insertion of mNG-mAID at the C-terminus. (**B**) Western blot from total bradyzoite lysates of conditional KD of TgPROP2-mAID with IAA or vehicle control for 72 h. Serial dilution of vehicle treated bradyzoites were used for quantification of protein levels, with relative percentages denoted above each lane. (**C**) Immunofluorescence assay of *in vitro* TgPROP2-mAID differentiated cysts treated with IAA or vehicle control for 72 h for conditional knockdown (KD). n = 3 biological replicates, representative images shown. Scale bars, 10 μm. (**D**) Autolysosome staining (CytoID) of *in vitro* TgPROP2-mAID, TgATG9-mAID, or TIR1 differentiated cysts treated with LHVS and IAA or vehicle control for 72 h. TIR1 treated with DMSO (top left quadrant) shown as control for the necessity of LHVS in the accumulation of autolysosomes. n = 3 biological replicates, representative images shown. Scale bars, 10 μm. (**E**) Quantification of (D) as number of CytoID puncta per cyst. Each point represents a single cyst with different shapes corresponding to each biological replicate. Bars represent mean ± SD. Statistical analysis was done using ordinary two-way ANOVA with Tukey’s multiple comparisons with p-values denoted above bars (n=3 biological replicates, ≥10 cysts quantified per replicate).

### TgPROP1 plays a key role in bradyzoite autophagy

Since our findings suggest that TgPROP2 is not required for autophagy, we hypothesized that TgPROP1 may be the critical protein for bradyzoite autophagy among the two Atg18/WIPI paralogs in *T. gondii*. TgPROP1 is not fitness-conferring in tachyzoites based on a genome-wide CRISPR-Cas9 screen (30), enabling direct genetic disruption. We generated TgPROP1 knockout (KO) parasites in a ME49Δku80Δhxg strain background (termed WT hereafter) using CRISPR-Cas9 by replacing the entire genomic sequence with the hypoxanthine-xanthine-guanine phosphoribosyl transferase (HXGPRT) selectable marker (35). Thereafter, we genetically complemented the TgPROP1 KO strain by reintroducing the genomic sequence of TgPROP1 harboring a silent mutation in a HindIII restriction site to distinguish it from WT (Figure 2A).

**Figure 2.**
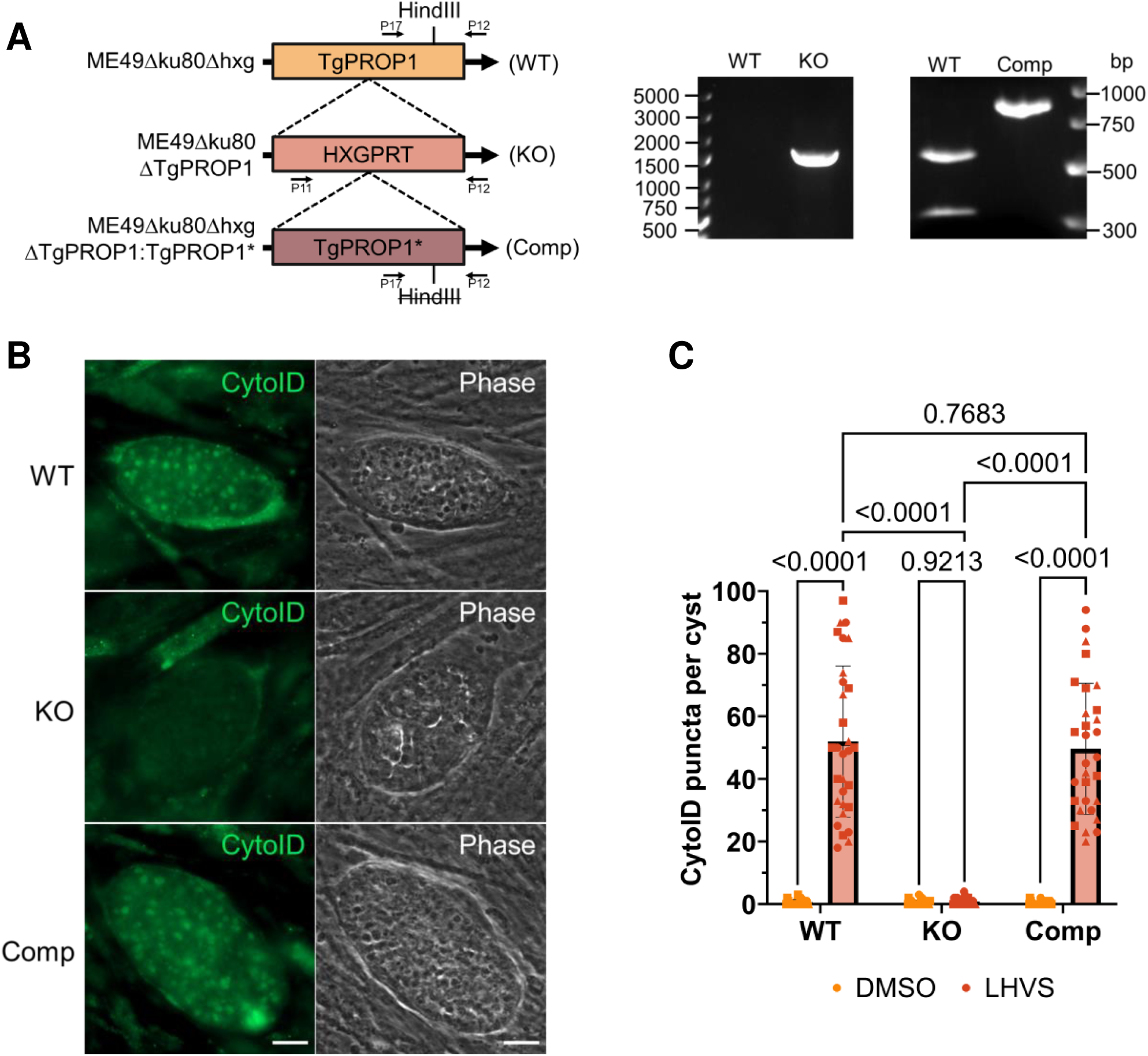
TgPROP1 plays a key role in bradyzoite autophagy. (**A**) Genetic ablation of TgPROP1 was achieved with CRISPR-Cas9 and homology-directed insertion of the hypoxanthine-xanthine-guanine phosphoribosyl transferase (HXGPRT) gene into the TgPROP1 locus (KO). Complementation of the KO was generated by CRISPR-Cas9 and homology-directed insertion of the genomic DNA sequence of TgPROP1 with a silent mutation in a HindIII restriction site at the C-terminus (Comp). PCR validation for the insertion of the HXGPRT gene in the TgPROP1 gene in the KO clone isolated. PCR amplified fragments by (indicated by arrows) from WT and Comp with HindIII restriction digestion. (**B**) Autolysosome staining (CytoID) of *in vitro* WT, TgPROP1 KO, or TgPROP1 Comp differentiated cysts treated with LHVS for 72 h. n = 3 biological replicates, representative images shown. Scale bars, 10 μm. (**C**) Quantification of (B) as number of CytoID puncta per cyst. Each point represents a single cyst with different shapes corresponding to each biological replicate. Bars represent mean ± SD. Statistical analysis was done using ordinary two-way ANOVA with Tukey’s multiple comparisons with p-values denoted above bars (n=3 biological replicates, ≥10 cysts quantified per replicate).

We assessed autophagic function of each strain (WT, KO, Comp) by differentiating *in vitro* bradyzoites for 7 days followed by treatment with LHVS or DMSO for 3 days prior to CytoID staining. CytoID-positive autolysosomes were observed in both WT and TgPROP1 Comp bradyzoites. In contrast, TgPROP1 KO bradyzoites were unable to accumulate CytoID-positive structures, indicating an impairment in autophagy (Figure 2B-C). These results provide evidence for the key role of TgPROP1 in bradyzoite autophagy and suggest that TgPROP1 performs a non-redundant function in this pathway.

### TgPROP1 is critical for T. gondii persistence in a mouse model of chronic infection

To determine whether TgPROP1 is necessary for the persistence of *T. gondii in vivo*, we infected mice with WT, TgPROP1 KO, or TgPROP1 Comp parasites. At 5 weeks post-infection, brains from infected mice were collected and cyst burdens enumerated. We observed a ∼40 fold and ∼20-fold decreased cyst burden in the brains of mice infected with TgPROP1 KO parasites compared to WT or the TgPROP1 Comp, respectively (Figure 3A). To visually assess the health of *ex vivo* cysts, we examined cysts from mouse brains via light microscopy. WT and TgPROP1 Comp cysts generally had bright, smooth, and distinctive cyst walls surrounding numerous bradyzoites with minimal gaps within each cyst. In contrast, TgPROP1 KO bradyzoites appeared bloated, mottled, and with increased gaps between parasites, surrounded by a less defined cyst wall (Figure 3B). We isolated bradyzoites from a subset of the brains harvested (3 per strain) using pepsin-treatment and evaluated their *ex vivo* viability via plaque assay normalized to the number of parasite genomes by qPCR (36). We observed no plaques after 14 days from *ex vivo* TgPROP1 KO parasites, while WT and TgPROP1 Comp parasites were able to grow and produce plaques (Figure 3C-D). These findings confirm the importance of TgPROP1 and bradyzoite autophagy to the overall health and survival of *T. gondii* during chronic infection.

**Figure 3.**
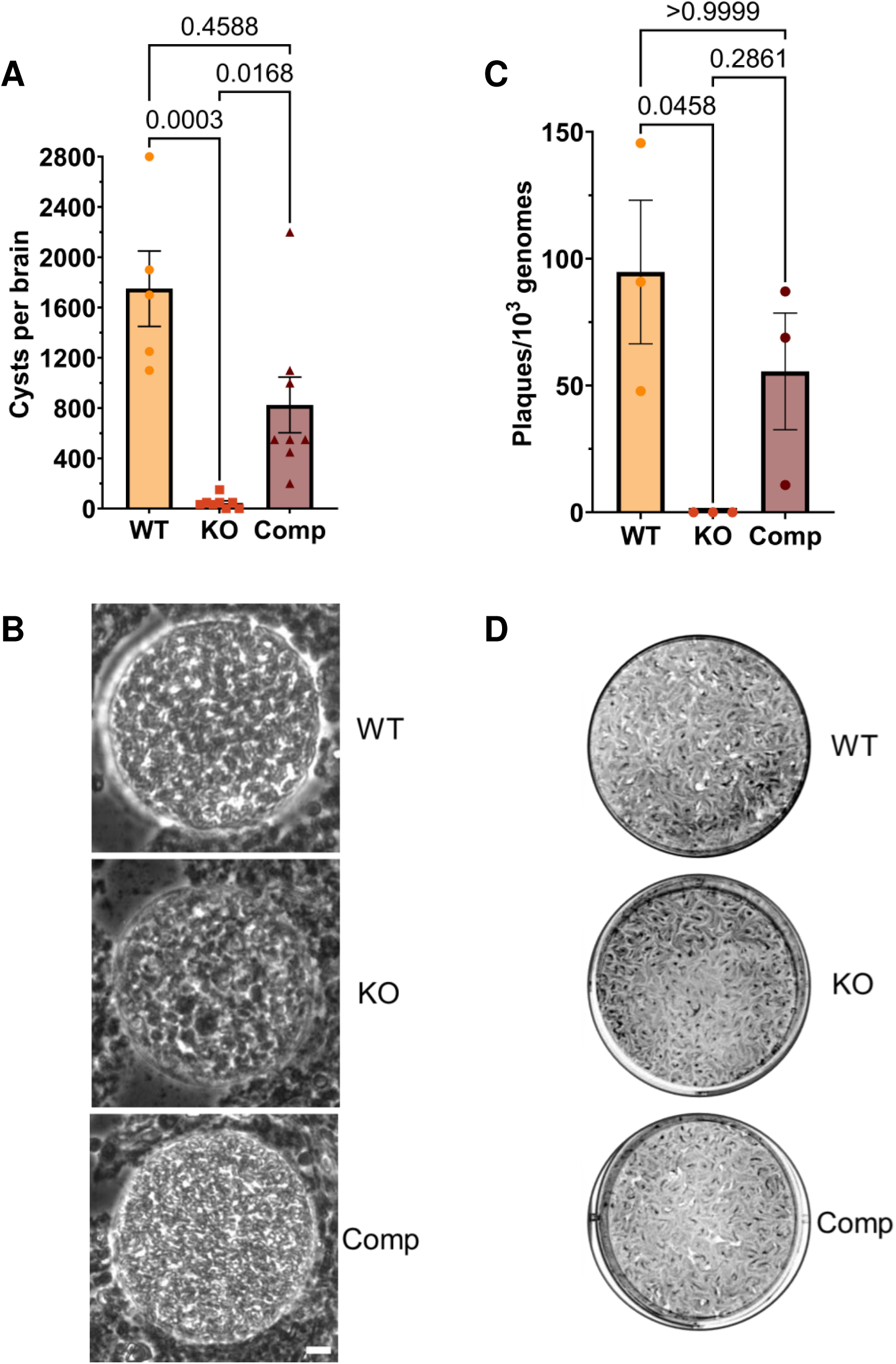
TgPROP1 is required for *T. gondii* persistence and viability during chronic infection. (**A**) Cyst burdens per brain quantified from chronically infected mice at 5-weeks post-infection with WT, KO, or Comp parasites. Brain samples were blinded and counted in triplicate using light microscopy. Bars represent mean ± SD. Statistical analysis was done using the Kruskal-Wallis one-way ANOVA test with p-values denoted above bars. (**B**) Representative images of ex vivo cysts from infected mouse brains under light microscopy for WT, KO, or Comp. Scale bars, 10 μm. (**C**) Viability of *ex vivo* bradyzoites harvested from infected mouse brains (3 representative brains per strain) from (A). Viability is reported as number of plaques normalized to the number of parasite genomes. Bars indicate mean ± SD. Statistical analysis was done using the Kruskal-Wallis one-way ANOVA test with p-values denoted above bars. (**D**) Representative wells from plaque assay from (C) for WT, KO, or Comp. (n = 3 biological replicates).

## Discussion

Autophagosome biogenesis and maturation requires recruitment of multiple proteins, including protein scaffolds that serve as platforms for coordinated assembly of core autophagy machinery (37, 38). A key event in phagophore development is the generation of PtdIns3P on autophagic membranes that recruit PtdIns3P-binding proteins which promote reversible interactions among other autophagy proteins (7). The PROPPIN family proteins (Atg18, Atg21, and Hsv2) in yeast and WIPIs 1-4 in mammalian cells are important for this step. Characterization of their functions has revealed some compensatory redundancies among the paralogs in both yeast and mammalian cells (10–13, 28). While much less is known about the early steps of autophagy within the protozoan parasite *T. gondii*, two paralogs of Atg18/WIPI (TgPROP1 and TgPROP2) have been discovered and partially characterized to localize to autophagosomes upon cellular stress (26). However, their specific functional importance to *T. gondii* autophagy was unclear. In this study, we provide evidence that suggests TgPROP1 performs a non-redundant function in chronic stage *T. gondii*.

Since autophagy is not an essential pathway for normal growth of acute stage parasites (tachyzoites) but is required for chronic stage (bradyzoite) survival, we investigated the individual contributions of TgPROP1 or TgPROP2 to bradyzoite autophagy (15, 18–20). While conditional knockdown of TgPROP2 did not impair autophagy based on CytoID staining, the TgPROP1 KO failed to accumulate autolysosomes upon inhibition of proteolytic turnover within the PLVAC. Although we confirmed marked knockdown of TgPROP2 by western blotting and immunofluorescence, we cannot rule out that residual amounts of TgPROP2 could have retained autophagy function. Nonetheless, that disrupting TgPROP1 resulted in a near complete absence of CytoID puncta indicates that TgPROP2 cannot compensate for, and is therefore, non-redundant with TgPROP1. This phenotype is like that observed for the TgATG9 KO, for which only a single homolog exists in *T. gondii* (19). These results align with prior sequence comparisons showing that the sequence similarity to Atg18 is higher for TgPROP1 (34%) than it is for TgPROP2 (29%) (26). It is well recognized that many of the autophagy proteins found in *T. gondii* have been repurposed for a non-canonical role in apicoplast homeostasis, including TgPROP2 and TgATG8 (23–25, 27, 39–42). Our results suggest that TgPROP2 plays little to no role in canonical autophagy and that its PtdIns3P-binding capabilities may be solely for the coordination of TgATG8 lipidation to apicoplast membranes rather than autophagic membranes (26, 27, 43). Future studies should aim to resolve the interacting network of TgPROP1 which may reveal new candidates distinctly dedicated to canonical autophagy in *T. gondii*.

We confirmed the necessity of TgPROP1 and autophagy for bradyzoite survival using a mouse model of chronic *T. gondii* infection. After 5 weeks, brains from mice infected with TgPROP1 KO parasites had a significant reduction in cyst burdens compared to WT or Comp strains. Bradyzoites derived from TgPROP1 KO infected brains were non-viable, failing to produce plaques after transition to normal growth conditions in tissue culture. We visually inspected the morphology of *ex vivo* cysts by light microscopy and observed that TgPROP1 KO cysts exhibited an overall unhealthy, bloated, and mottled appearance. These observations suggest that TgPROP1 KO cysts might not be able to undergo recrudescence, an important feature in toxoplasmosis pathology within host organs (44–47). Due to its critical importance in chronic infection, *T. gondii* autophagy remains a pathway worthwhile of future studies that could lead to pragmatic gains.

Overall, our research has unveiled a unique, non-redundant role of TgPROP1 in *T. gondii* autophagy, substantiated by both *in vitro* and *in vivo* studies. While numerous aspects of the function and regulation of the autophagy pathway in this divergent eukaryote remain to be elucidated, our study highlights the pivotal role of autophagy in bradyzoite survival. These findings have direct implications for developing innovative strategies to combat *T. gondii* chronic infection.

## Materials and Methods

### Parasite and host cell culture

*T. gondii* parasites were cultured in human foreskin fibroblasts (HFFs, Hs27) obtained from the American Type Culture Collection (ATCC-CRL-1634). Tachyzoite growth was maintained at 37°C under 5% CO_2_ in D10 medium, composed of DMEM (Fisher Scientific, 10-013 cv), 10% heat-inactivated bovine calf serum (Cytiva, SH30087.03), 2 mM L-glutamine (Corning, 25-005-Cl), and 50 U/mL of penicillin-streptomycin (Gibco, 15070063). For bradyzoite differentiation, tachyzoites were mechanically lysed from infected HFF monolayers via scraping and syringing (20- and 25-gauge needles) then allowed to invade fresh monolayers of HFFs for 24 h at 37°C under 5% CO_2_ in D10 medium. The next day, D10 medium was replaced with alkaline-stress medium consisting of RPMI-1640 without NaHCO_3_ (Cytiva, SH30011.02) supplemented with 3% heat-inactivated fetal bovine serum (Cytiva, SH30396.03), 50 mM HEPES (Sigma-Aldrich, H3375), 50 U/mL of penicillin-streptomycin and adjusted to pH 8.2-8.3 with NaOH. Bradyzoites were cultured at 37°C under ambient CO_2_ with daily medium changes to maintain alkaline pH (48).

### T. gondii transfection

Tachyzoites were mechanically lysed from fully infected HFF monolayers via scraping and syringing (20- and 25-gauge needles). Parasites were pelleted and resuspended in cytomix transfection buffer (2 mM EDTA, 120 mM KCl, 0.15 mM CaCl_2_, 10 mM K_2_HPO_4_/KH_2_PO_4_, 25 mM HEPES, 5 mM MgCl_2_, pH 7.6). For each transfection, guide RNA plasmids, homology-direct repair templates, or linearized plasmids were precipitated with ethanol and resuspended in cytomix buffer. The DNA mixture was combined with pelleted parasites prior to electroporation in 0.4-cm cuvettes (Bio-Rad, 1652081), utilizing the GenePulser Xcell with PC and CE modules (Bio-Rad, 1652660), and configured with the following parameters: 2400 V voltage, 25 μF capacitance, 50 Ω resistance.

### T. gondii strain generation

Primers and oligos were synthesized either by IDT or Sigma-Aldrich. All guide RNAs were generated by substituting the original 20 base pair guide RNA sequence on the plasmid pCas9/sgRNA/Bleo (49) with the desired 20 base pair guide RNA sequence, using the Q5® Site-Directed Mutagenesis Kit (New England Biolabs, E0554S). Homology-directed repair templates were PCR amplified using the CloneAmp™ HiFi PCR Premix (Takara Bio, 639298). Guide RNA and repair template sequences are listed in Table S1.

### Generation of the TgPROP2-mAID T. gondii strain

Starting with the TIR1-expressing ME49 *T. gondii* strain (ME49/TIR1) (20, 50), TgPROP2 was endogenously tagged at the C terminus to generate ME49/TIR1/TgPROP2-mAID (TgPROP2-mAID) parasites. A guide RNA targeting the C terminus of TgPROP2 near the stop codon was generated using oligos P1/P2. Homology-directed repair template was generated using oligos P3/P4 for PCR amplification of pGL015 (51) containing the Xten linker, V5 epitope (Invitrogen, 37-7500), mNeonGreen, minimal AID, and Ty epitope. ME49/TIR1 parasites were co-transfected with 100 µg guide RNA and 50 µg repair template. At 48 h after transfection, positively transfected parasites were selected using 5 µg/mL phleomycin (Invitrogen, NC9198593) prior to isolating clones by limiting dilution. Individual parasite clones were validated by PCR amplification to confirm presence of the mAID tag using P5/P6, live imaging for mNeonGreen fluorescence, and immunofluorescence.

### Generation of the TgPROP1 KO T. gondii strain

Starting with the ME49*Δku80Δhxgprt T. gondii* strain (52), the entire genomic DNA sequence of TgPROP1 was replaced with the hypoxanthine-xanthine-guanine phosphoribosyl transferase (HXGPRT) gene to generate ME49*Δku80ΔTgPROP1* (TgPROP1 KO) parasites. Two guide RNAs targeting the N terminus and C terminus of TgPROP1 near the start or stop codons, respectively, were generated using oligos P7/P2 and P8/P2. Homology-directed repair template was generated using oligos P9/P10 for PCR amplification of the HXGPRT selectable marker (53). ME49*Δku80Δhxgprt* parasites were co-transfected with 50 µg of each guide RNA and 50 µg repair template. At 24 h after transfection, positively transfected parasites were selected using 25 µg/mL mycophenolic acid and 50 µg/mL xanthine (Sigma-Aldrich, M3536 and X3627) for 5 days prior to isolating clones by limiting dilution. Individual parasite clones were validated by PCR amplification to confirm presence of the HXGPRT insertion using P11/P12.

### Generation of the TgPROP1 Comp T. gondii strain

Starting with the ME49*Δku80ΔTgPROP1* (TgPROP1 KO) *T. gondii* strain, the HXGPRT selectable marker was replaced with the full TgPROP1 gene to generate ME49*Δku80ΔTgPROP1:TgPROP1\** (TgPROP1 Comp) parasites. Two guide RNAs targeting the N terminus and C terminus of TgPROP1 (replaced by HXGPRT) near the start or stop codons, respectively, were generated using oligos P13/P2 and P14/P2. A homology-directed repair template was generated using oligos P15/P16 for PCR amplification of a plasmid synthesized to contain the full TgPROP1 genomic sequence with a silent mutation at a HindIII restriction site near the C terminus (GeneUniversal, Table S1). TgPROP1 KO parasites were co-transfected with 50 µg of each guide RNA and 50 µg repair template. At 48 h after transfection, positively transfected parasites were selected using 5 µg/mL phleomycin (Invitrogen, NC9198593) prior to isolating clones by limiting dilution. Individual parasite clones were validated by PCR amplification to confirm presence of the TgPROP1 gene using P17/P12 and to confirm the silent mutation in the HindIII site, the small, amplified fragment of the C terminus of the TgPROP1 gene from ME49*Δku80Δhxgprt* (WT) and TgPROP1 complemented parasites were subjected to HindIII restriction digestion (New England Biolabs, R3104S).

### Western blotting

For validation of TgPROP2 knockdown by western blotting, TgPROP2-mAID parasites were differentiated into bradyzoites for 7 days grown in alkaline-stress medium (ambient CO_2_) with daily media changes. On day 7, bradyzoites were treated with 500 µM IAA (Sigma-Aldrich, I2886) or ethanol vehicle control for 24 h. Bradyzoites were harvested via scraping, syringing (20- and 25-gauge needles), pepsin treatment (0.026% pepsin [Sigma-Aldrich, P7000] in 170 mM NaCl and 60 mM HCl, final concentration), and filtration. Bradyzoites were enumerated and lysed with RIPA buffer (Thermo Scientific, 89900) supplemented with cOmplete Mini Protease Inhibitors cocktail (Roche, 11836153001) for 15 min at 4°C with gentle rocking. Lysates were centrifuged at 4°C for 10 min at 20,000 x g. Lysates were supplemented to a final concentration of 1X SDS-PAGE buffer and 2% β-mercaptoethanol. The final concentration was ∼1×10^7^ bradyzoites per 25 µL and designated as 100% for loading.

Lysates were subjected to SDS-PAGE using a gradient 4-12% NuPAGE™ Bis-Tris gel (Invitrogen, NP0321) and transferred onto 0.45-µm nitrocellulose membranes (Bio-Rad, 1620115) with Trans-Blot® SD semi-dry transfer cell (Bio-Rad, 1703940) for 45 min at 18 V at room temperature. Following transfer, membranes were blocked with 5% milk in phosphate-buffered saline (PBS [Gibco, 21600010])-T (PBS with 0.05% Triton X-114 [Sigma-Aldrich, X114] and 0.05% Tween-20 [Fisher Scientific, 170-6531]) for 30 min at room temperature. Primary antibodies were diluted in 1% milk in PBS-T and applied to membranes overnight at 4°C. Primary antibodies used include mouse anti-mNeonGreen (Chromotek, 32F6; 1:1000) and rabbit anti-TgActin (Sibley lab, Washington University in St. Louis; 1:20,000). After primary antibody incubation, membranes were washed 3 times with PBS-T before incubation with HRP-conjugated secondary antibodies (Jackson ImmunoResearch Laboratories, 115-035-146; 1:5000) for 1 h at room temperature. Proteins were detected using SuperSignal™ West Pico PLUS Chemiluminescent Substrate or Femto Maximum Sensitivity Substrate (Thermo Fisher, 1863096 or 34095). The Syngene PXi6 imaging system with Genesys (v1.8.2.0) software was used to detect signals.

### Immunofluorescence

*T. gondii* ME49/TIR1 or ME49/TIR1/TgPROP2-mAID tachyzoites were differentiated into bradyzoites in 6-well tissue culture plates with 22 mm x 22 mm No. 1.5 coverslips (Globe Scientific, 1404-15) for 7 days. Bradyzoites were fixed with 4% paraformaldehyde, permeabilized with 0.1% Triton X-100 in PBS for 10 min and blocked with 10% FBS in PBS (with 0.01% Triton X-100) for 30 min. Primary and secondary antibodies, along with other staining reagents, were diluted in wash buffer (1% FBS, 1% normal goat serum [Gibco, 16210072], and 0.01% Triton X-100 in PBS). All antibody incubations were performed at room temperature for 1 h. The primary antibody used was mouse anti-mNeonGreen (Chromotek, 32F6; 1:1000). The secondary antibody used was goat anti-mouse Alexa Fluor 488 (Invitrogen, A28175; 1:1000). Coverslips were mounted using ProLong™ Gold Antifade Mountant (Thermo Scientific, P36930). Images were taken on a Zeiss Axio Observer Z1 inverted microscope at 63× and analyzed using Zen 3.7 blue edition software.

### CytoID staining of autolysosomes

*T. gondii* tachyzoites were differentiated into bradyzoites in 6-well tissue culture plates with 22 mm x 22 mm No. 1.5 coverslips (Globe Scientific, 1404-15) for 7 days then treated with 1 µM LHVS (Sigma-Aldrich, SML2857) or equal volume DMSO for 3 days. For knockdown of TgPROP2-mAID, TgATG9-mAID, and ME49/TIR1 controls, bradyzoites were also concurrently treated with 500 µM IAA or equal volume ethanol for 3 days. The CytoID Autophagy Detection Kit 2.0 (Enzo, ENZ-KIT175) was used to stain autolysosomes within live bradyzoites for 45 min prior to fixation with 4% paraformaldehyde, following the manufacturer’s instructions. Fixed coverslips were mounted using ProLong™ Gold Antifade Mountant (Thermo Scientific, P36930). Images were taken on a Zeiss Axio Observer Z1 inverted microscope at 63× and analyzed using Zen 3.7 blue edition software. For quantification of the number of CytoID puncta within each cyst, CytoID-positive puncta were enumerated from at least 10 cysts per biological replicate, from 3 biological replicates total. All images were coded prior to quantification to blind the experimenter during quantification.

### Mouse T. gondii chronic infection

Eight-week-old female CBA/J mice (Jackson Laboratory, 000656) were randomly assigned to groups and infected intra-peritoneally (i.p.) with 500 tachyzoites of ME49*Δku80* (n = 10 mice), ME49*Δku80ΔTgPROP1* (n = 8 mice), or ME49*Δku80ΔTgPROP1:TgPROP1\** (n = 8 mice) strains. At 5 weeks post-infection, mice were humanely euthanized and cyst burdens in the brains were assessed. Mouse brains were individually homogenized with scissors followed by multiple passages through a 21G syringe needle in a final volume of 1 mL in PBS. All samples were blinded prior to enumeration of cysts. Three 10 μL samples (30 μL total) of brain homogenates per infected mouse were analyzed by light microscopy to enumerate cysts and the number of cysts per brain determined by scaled calculation to the homogenate volume. Animal studies described here adhere to a protocol approved by the Committee on the Use and Care of Animals of the University of Michigan.

### Ex vivo bradyzoite viability assay

*Ex vivo* brains from chronically infected mice were homogenized in 1 mL of sterile Hanks’ buffered salt solution (Gibco, 14175103), and bradyzoites were subsequently harvested using pepsin (0.026% pepsin in 170 mM NaCl and 60 mM HCl, final concentration) treatment (Sigma-Aldrich, P6887) (36). The viability of the purified *ex vivo* bradyzoites was assessed by combining plaque assay and qPCR normalization of parasite genome numbers. Equal volumes of bradyzoites were added in triplicate to fresh 6-well tissue culture plates containing confluent HFFs in D10 medium. Bradyzoite-derived plaques were allowed to form undisturbed for 14 days, grown at 37°C under 5% CO_2_. Plaques were counted using a light microscope and plates were stained with crystal violet fix solution (0.2% of crystal violet and 70% of EtOH) for 20 min at room temperature. Genomic DNA from pepsin-treated bradyzoites was obtained with the DNeasy Blood and Tissue Kit (Qiagen, 69506). qPCR was performed using 2 µL of genomic DNA in SsoAdvancedTM Universal SYBR® Green Supermix (Bio-Rad, 172-5271) and primers for a 529-base pair repetitive element of *T. gondii* (forward AGGAGAGATATCAGGACTGTAG; reverse GCGTCGTCTCGTCTAGATCG) (54). The qPCR reactions were performed with the CFX96 Touch Real-Time PCR Detection System (Bio-Rad) using the following parameters: 3 min at 98°C, and 40 cycles of 15 s at 98°C, 30 s at 58.5°C, and 30 s at 72°C. A standard curve was built with 6.4, 32, 160, 800, 4000, 2000 parasite genomes. The number of plaques was normalized to the calculated number of genomes present in the inoculating samples.

### Statistical analysis

Data were analyzed using GraphPad prism. For each data set, outliers were identified and removed using ROUT with a *Q* value of 0.1%. Data were then tested for normality and equal variance. Student’s *t*-test or one-way analysis of variance (ANOVA) was performed for normally distributed data with equal or assumed equal variance, when appropriate. If the data failed one or both tests, a Kruskal-Wallis test was performed. Specific details of each test are described in the corresponding figure legends.

## Abbreviations

PROPPIN: β-propeller that bind phosphoinositides
PtdIns3P: phosphatidylinositol-3-phosphate
PtdIns3K: phosphatidylinositol 3-kinase
IAA: indole-3-acetic acid
mAID: minimal auxin-inducible degron
PLVAC: plant-like vacuolar compartment

## Acknowledgements

We thank the Sibley lab at Washington University in St. Louis for the TgActin antibody.

